# The conserved metalloprotease invadolysin is present in invertebrate haemolymph and vertebrate blood

**DOI:** 10.1101/612127

**Authors:** Kanishk Abhinav, Linda Feng, Emma Morrison, Yunshin Jung, James Dear, Satoru Takahashi, Margarete M. S. Heck

## Abstract

We identified invadolysin, a novel essential metalloprotease, for functions in chromosome structure, cell proliferation and migration. Invadolysin also plays an important metabolic role in insulin signaling and is the only protease known to localise to lipid droplets, the main lipid storage organelle in the cell. *In silico* examination of the protein sequence of invadolysin predicts not only protease and lipase catalytic motifs, but also post-translational modifications and the secretion of invadolysin. Here we show that the protease motif of invadolysin is important for its role in lipid accumulation, but not in glycogen accumulation. The lipase motif does not appear to be functionally important for accumulation of lipids or glycogen. Post-translational modifications likely contribute to modulating the level, localisation or activity of invadolysin. We identified a secreted form of invadolysin in the soluble fraction of invertebrate hemolymph (where we observe sexually dimorphic forms) and also vertebrate plasma, including in the extracellular vesicle fraction. Biochemical analysis for various post-translational modifications demonstrated that secreted invadolysin is both N-and O-glycosylated, but not apparently GPI-linked. The discovery of invadolysin in the extracellular milieu suggests a role for invadolysin in normal organismal physiology.

**Summary Statement:** In this study, we show that the conserved metalloprotease invadolysin is present in invertebrate hemolymph and vertebrate blood, suggesting the protein may function in organismal physiology.

## Introduction

Proteases perform a wide array of functions in normal physiology ranging from cell proliferation, differentiation and death - to digestion, blood coagulation and complement pathway activation (Lecker *et al*., 2006; Vandenabeele *et al.*, 2005; Walsh and Ahmad, 2002; Werb *et al.*, 1999). *In silico* analyses of metazoan genomes have identified more than 500 proteases and inhibitors accounting for approximately 2-5% of total gene number (Puente *et al.*, 2003; Turk, 2006). With such a high percentage of the genome dedicated to protein turnover, it is somewhat surprising that to date, only a small fraction of these enzymes have been thoroughly investigated. Therefore, the characterisation of novel proteases is an important area of investigation, improving our understanding of the role of proteases in normal physiology and disease pathophysiology.

The ability to perform a wide variety of functions, coupled with the modulation of enzymatic activity, make proteases attractive drug targets. ACE (Angiotensin-Converting Enzyme) inhibitors are widely used for treating hypertension, myocardial infarction and renal failure (Wong *et al*., 2004). HIV (Human Immunodeficiency Virus) protease inhibitors have been successfully used to treat HIV-infected patients (Flexner, 1998). On the other hand, development of MMP inhibitors for treatment of connective tissue diseases failed during clinical trials, due largely to off-target effects (Cathcart and Cao, 2015). These results further signify the importance of a more thorough investigation of proteases and their activities to improve the development of protease-based therapies.

Invadolysin plays an important role in the cell cycle, cell migration and the maintenance of normal chromosome structure. Crucially, the gene is essential for life in *Drosophila* and plays analogous roles in zebrafish (McHugh *et al*., 2004; Rao *et al*., 2015)(Vass and Heck, 2013). Invadolysin has a conserved metalloprotease motif (HEXXH) and is the only member of the single-gene M8 family of metalloproteases in metazoa (McHugh *et al.*, 2004) – the prototype of this family being the Leishmanolysin/GP63 protease from *Leishmania*. To date, invadolysin is the only protease shown to localise to lipid droplets, the primary lipid storage organelle of the cell (Cobbe *et al.*, 2009). More recent studies have identified key roles for invadolysin in metabolism including insulin signaling, lipid accumulation and the maintenance of normal mitochondrial function (Chang *et al*., 2016; Di Cara *et al.*, 2013). Though a fairly conserved lipase motif lies downstream of the highly conserved protease motif, neither of these motifs has been thoroughly examined in invadolysin’s function. Are the protease and lipase motifs functional and do they contribute to invadolysin’s activity?

Protease activity may be regulated by various strategies such as irreversible activation of an inactive zymogen, reversible binding of cofactors, or exposure to different intra-or extra-cellular millieus (Twining, 1994). Physical subcellular compartmentalisation is also utilised to regulate protease activity (Brix *et al.*, 2013). Proteases may exist in soluble intracellular, membrane-bound, or extracellular forms. As proteases generally have numerous substrates, regulating substrate localization or accessibility will also serve to modulate activity (Schauperl *et al.*, 2015). Furthermore, post-translational modifications such as phosphorylation, N-or O-linked glycosylation or GPI-anchor addition may not only affect protein structure and stability, but also impact on substrate interaction and binding affinity (Goettig, 2016). All these factors further add to the complexity of the mechanisms regulating protease activity. In this study, we address the biosynthesis and post-translational modification of invadolysin.

The first evidence for a diffusible protease was demonstrated using a tadpole explant that was capable of degrading a collagen gel (Gross and Lapiere, 1962). A number of extracellular proteases have been discovered since. Proteases such as chymotrypsin, trypsin and carboxypeptidase (components of the digestive system) are responsible for hydrolysing proteins before absorption in the gastrointestinal tract (Szmola *et al*., 2011). Extracellular proteases play important roles in angiogenesis, tissue remodelling and wound healing (Birkedal-Hansen *et al.*, 1993; Verma and Hansch, 2007). The ADAMTSs (A Disintegrin And Metalloproteinase with ThromboSpondin motifs) family of extracellular metalloproteases are important for angiogenesis (Rodríguez-Manzaneque *et al.*, 2015), whereas secreted metalloproteases such as MMPs (Matrix MetalloProteinase) play vital roles in extracellular matrix remodelling (Birkedal-Hansen *et al.*, 1993). ADAMTSs and MMPs, each represented by complex multi-gene families, are some of the better-characterised secreted proteases. However, functional redundancy of other family members often complicates interpretation of phenotypic disruption.

We previously demonstrated that invadolysin plays a crucial role in metabolism and energy storage in *Drosophila* (Chang *et al.*, 2016; Cobbe *et al*., 2009). We set about further examining these functions using a number of approaches. We generated transgenic fly lines that expressed either wild type, protease-or lipase-dead forms of invadolysin and compared lipid and glycogen accumulation amongst them. *In silico* analysis of the invadolysin sequence identified potential sites of post-translational modifications, which suggested not only phosphorylation, glycosylation and GPI-anchor addition, but also the secretion of invadolysin. This led to our discovery of invadolysin in the soluble fraction of both vertebrate blood and invertebrate hemolymph. While secreted invadolysin is glycosylated, it does not appear to have a GPI-anchor. A portion n of this secreted invadolysin is present in a human plasma fraction enriched for extracellular vesicles, suggesting additional roles in mediating communication between cells or tissues. Our present study opens new avenues of research into the physiological role(s) of extracellular invadolysin.

## Results

### *In silico* identification of conserved sequence features of invadolysin

Metalloproteases are generally zinc-dependent enzymes that have a conserved HEXXH (zincin) or HXXEH (inverzincin) metalloprotease motif (Gomis-Rüth, 2003). In addition to having the classical HEXXH zincin metalloprotease motif, invadolysin also has a third conserved histidine residue and a downstream Met-turn (Figure 1, red and green), placing invadolysin in the M8 subfamily of metalloproteases (McHugh *et al*., 2004). Leishmanolysin/GP63, a major surface protease of *Leishmania major* (though also found intra-and extra-cellularly), is the prototype for the M8 leishmanolysin subfamily of metalloproteases (Gomis-Rüth, 2003; McGwire et al., 2002). Invadolysin additionally has a conserved lipase (GXSXG) motif just downstream of the protease motif (Figure 1, purple). The lipase motif consists of two glycines and a serine where the serine is the catalytically-active residue (Wong and Schotz, 2002).

**Figure 1.**
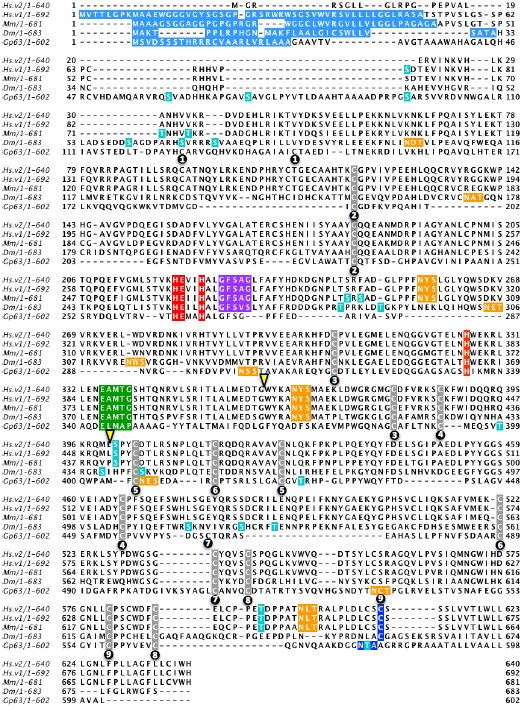
Multiple sequence alignment of invadolysin protein sequence from *Homo sapiens* (Hs.v2: variant with an alternative start site; Hs.v1: the more abundant invadolysin variant), *Mus musculus* (Mm), *Drosophila melanogaster* (Dm) with Gp63 (Leishmanolysin) using ClustalX. The schematic was generated using Jalview to highlight sequence features within the protein. Signal sequence was predicted using SignalP 4.1 Server (light blue), O-glycosylation sites were predicted using NetOGlyc 4.0 Server - DTU CBS (turquoise), N-glycosylation was predicted using NetNGlyc 1.0 Server - DTU CBS (orange) and GPI-anchor addition was predicted using big-PI Predictor (darker blue). The conserved metalloprotease motif and the third histidine residue are highlighted in red and the downstream Met-turn is highlighted in green. The conserved metalloprotease motif, third histidine and the downstream Met-turn are the key identifying features of metalloprotease that places invadolysin in the Leishmanolysin M8 subclass of metalloprotease. Adjacent to the conserved metalloprotease motif in invadolysin is a fairly conserved lipase motif (purple). Cysteine residues conserved between invadolysin and leishmanolysin are highlighted in grey and those that have been identified to form disulphide bonds in Leishmanolysin have been numbered (Schlagenhauf *et al.*, 1998). An alternatively spliced 37 amino acid exon (in human) is between the two yellow arrowheads.

Human invadolysin is represented by 4 different variants – two for each of two different N-terminal variants which vary by alternative splicing of a 37 amino acid exon (between yellow arrowheads). Invadolysin variant 1 is predicted to encode an N-terminal signal sequence when analysed by SignalP 4.1 (a signal sequence prediction server) (Figure 1, light blue) (Nielsen *et al.*, 1997; Petersen *et al.*, 2011). Variant 2 (not found in mouse or fly) is not predicted to encode an N-terminal signal sequence. Synthesis of variant 2 is dependent on the use of an alternative translation start site (Cobbe *et al*., 2009). Though variant 2 can be detected by RT-PCR, it is not as prevalent as variant 1, and under what conditions it is translated is currently under investigation. We have identified the expression of variant 2 during the later stages of *in vitro* adipogenesis (Chang *et al.*, 2016). *in silico* analysis of invadolysin open reading frames from different species thus suggests the presence of a signal sequence that could target the translation of invadolysin to the secretory pathway.

The classical or conventional pathway of protein secretion utilises an N-terminal signal sequence to target the nascent protein to the endoplasmic reticulum and subsequently to the Golgi apparatus (Lippincott-Schwartz *et al.*, 2000). The protein may then translocate from the Golgi apparatus to the cell surface or be secreted via extracellular vesicles (Bendtsen *et al.*, 2004; Lippincott-Schwartz *et al.*, 2000). Proteins frequently undergo assorted post-translational modifications during their transport within the secretory pathway. N-glycosylation and C-terminal GPI-anchor addition occur within the endoplasmic reticulum (Aebi, 2013; Eisenhaber *et al.*, 2001), while O-glycosylation occurs in the Golgi apparatus (Spiro, 2002). Big-PI Predictor (Eisenhaber *et al.*, 1999) predicted the presence of a GPI-anchor site near the C-terminus of invadolysin (Figure 1, dark blue). A GPI-addition could anchor invadolysin to the plasma membrane (Fujita and Kinoshita, 2012; Orlean and Menon, 2007). The presence of several N-glycosylation (orange) and O-glycosylation (turquoise) sites are also predicted for invadolysin, (NetNGlyc 1.0) (Gupta and Brunak, 2002) and (NetOGlyc 3.1) (Steentoft *et al.*, 2013) respectively. Biochemical analysis of these predicted motifs in invadolysin is addressed below.

As highlighted in our identification of invadolysin (McHugh *et al.*, 2004), the higher eukaryotic forms of invadolysin all contain distinct regions of sequence that are not present in leishmanolysin (Figure 1, visible as gaps in the bottom row of the alignment). In spite of this, 9 pairs of cysteines are conserved in spacing and position (Figure 1, grey), suggesting that the structural ‘core’ of invadolysin may resemble that of leishmanolysin. The numbered black circles represent which cysteines are disulphide-bonded with one another within the leishmanolysin crystal structure (Schlagenhauf *et al.*, 1998).

### Proteolytic activity of invadolysin is important for its role in lipid accumulation

Our previous studies suggested a crucial role for invadolysin in energy storage. *invadolysin* mutant third instar larvae have reduced fat body thickness and cellular cross-sectional area (Cobbe et al., 2009). Critically, *invadolysin* mutants also have reduced triglyceride and glycogen levels (Chang *et al.*, 2016; McHugh *et al.*, 2004). To further analyse the role of invadolysin’s conserved protease and lipase catalytic motifs in lipid and glycogen storage, transgenic *Drosophila* strains with mutated motifs were generated by site-directed mutagenesis. A protease dead (E258A) form of invadolysin was generated by mutating the glutamic acid residue within the protease motif, whilst a lipase dead (S266A) form was generated by mutating the serine residue within the lipase motif (Figure 2A). Mutant versions of invadolysin were placed under the control of a UAS promoter (Brand and Perrimon, 1993).

**Figure 2.**
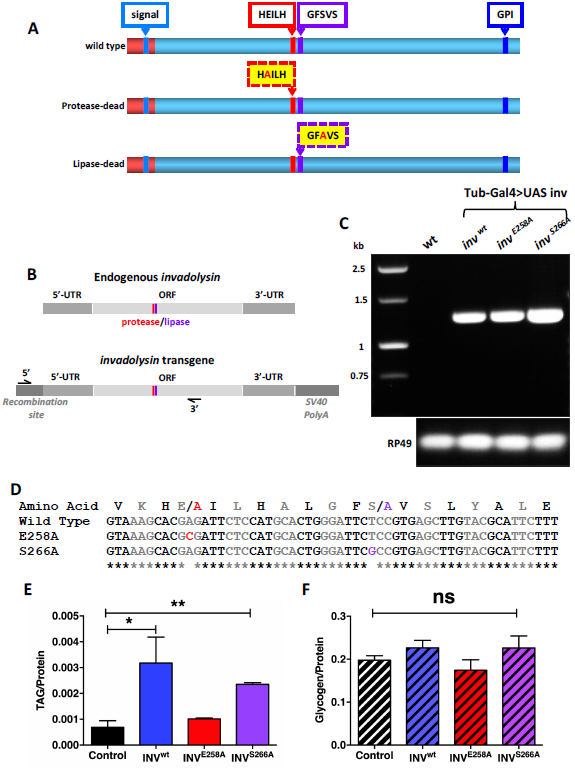
A) Schematic of wild type, protease-and lipase-dead versions of invadolysin highlighting the sites of the introduced mutations. B) Schematic of endogenous and transgenic invadolysin RNA illustrating the primers that selectively amplify transgenic invadolysin. C) RT-PCR showed that the invadolysin transgene can be expressed using a ubiquitous tubulin-Gal4 driver. D) Sequencing results from analysis of PCR amplicons confirmed that the transgenic flies have the induced mutations in the invadolysin transgenes. E) Triglyceride:protein ratio in flies overexpressing either wild type or mutant versions of invadolysin. Flies overexpressing wild type and the lipase-dead form of invadolysin had a significantly higher triglyceride:protein ratio than control flies, while flies overexpressing the protease-dead form of invadolysin failed to accumulate excess lipids, highlighting the importance of the metalloprotease motif in lipid accumulation. F) Glycogen:protein ratio in flies overexpressing either wild type or mutant versions of invadolysin. There was no significant difference in glycogen:protein ratio comparing the transgenic invadolysin strains with the control flies suggesting neither proteolytic nor lipolytic activity were directly important for glycogen accumulation.

*invadolysin* transgenes were integrated into a predetermined location within the genome (in this case, on the second chromosome) using the phiC31 integration system (Bischof *et al*., 2007). This system was exploited to minimise chromosomal positional effect on transgene expression which might affect levels or catalytic activity of the various invadolysin transgenes (Markstein *et al.*, 2008). *invadolysin* transgene expression was verified by RT-PCR using primers (diagrammed in Figure 2B) that selectively amplified the transgenic mRNA (Figure 2C, top panel). The PCR amplicons were subsequently sequenced to confirm the presence of the desired E258A and S266A mutations in the invadolysin transgenic mRNAs (Figure 2D).

Using the UAS-Gal4 system, we examined transgenic flies overexpressing wild type or mutant forms of invadolysin for triglyceride and glycogen levels. A tubulin-Gal4 driver was utilised to generate ubiquitous expression. Flies overexpressing wild type invadolysin accumulated significantly higher levels of triglyceride compared to control animals (Figure 2E). Flies overexpressing the *lipase*-dead form of invadolysin also accumulated higher levels of triglyceride, suggesting that this motif was not essential to accumulate increased triglyceride. On the other hand, the ability of flies to accumulate higher amounts of triglyceride was impaired upon overexpression of a *protease*-dead form of invadolysin. These data suggest intriguingly that invadolysin’s *proteolytic* activity is important for its role in lipid accumulation. Overexpression of any of the three invadolysin transgenes had no significant effect on glycogen accumulation (Figure 2F). We propose that the decreased glycogen level observed in *invadolysin* mutants is likely due to impaired insulin signaling or metabolism of glycogen reserves (Chang *et al.*, 2016).

### An extracellular form of invadolysin is present in *Drosophila* hemolymph

To examine if the predicted signal sequence in invadolysin led to secretion of the protein, *Drosophila* hemolymph was analysed by immunoblotting. Hemolymph is the invertebrate functional equivalent of vertebrate blood, and like blood in a vertebrate, is composed of a cellular component and soluble plasma (Wyatt *et al.*, 1956). The cellular component is comprised of hemocytes which include crystal cells, plasmatocytes, lamellocytes and precursor cells (Kurucz *et al.*, 2007). Immunoblotting of whole and fractionated hemolymph from adult male and female *Drosophila* identified that invadolysin was indeed present in soluble plasma, but intriguingly, that extracellular invadolysin differs in male (111 kDa) and female (46 kDa) flies (Figure 3B and C). We therefore examined male and female gonads by immunoblotting and observed that invadolysin similar in molecular weight to the secreted forms was present in testes and ovaries (Figure 3D and E). These results suggest testes and ovaries could be a source of secreted invadolysin, or alternatively, that these tissues import invadolysin from hemolymph.

**Figure 3.**
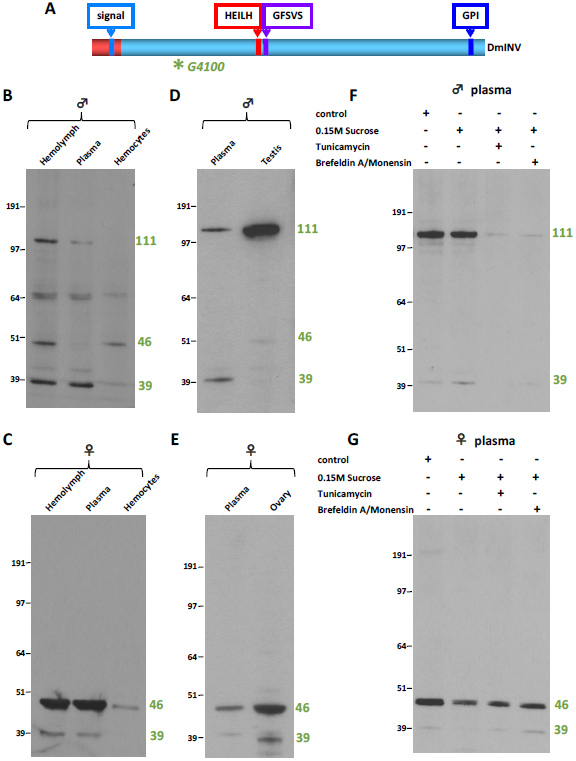
A) Schematic of *Drosophila* invadolysin with the G4100 antibody epitope highlighted. B) and C) Hemolymph fractions from male and female adult *Drosophila* respectively. A secreted form of invadolysin can be detected in the soluble hemolymph fraction of both males and females. D) and E) Invadolysin in gonads and soluble hemolymph in male and female *Drosophila* respectively show similar molecular weight forms. F) and G) Analysis of the effects of protein transport inhibitors on secretion of invadolysin in male and female *Drosophila* respectively. Feeding flies either Tunicamycin or Brefeldin A/Monensin resulted in decreased invadolysin in adult male plasma, but did not affect the level of invadolysin in adult female plasma.

As discussed previously, proteins encoding a signal sequence are targeted to the endoplasmic reticulum and subsequently to the Golgi apparatus and secretory vesicles (Bendtsen *et al.*, 2004). Several drugs can inhibit protein secretion by blocking the transport of vesicles along the secretory pathway. Brefeldin A disrupts protein transport from the endoplasmic reticulum to the Golgi apparatus by dissociating peripheral Golgi associated proteins. Monensin is a Na^+^ ionophore that disrupts transport within the Golgi apparatus (Helms and Rothman, 1992; Mollenhauer *et al.*, 1990). Tunicamycin inhibits N-glycosylation in the endoplasmic reticulum inducing endoplasmic reticulum stress which in turn inhibits protein secretion (Iwata *et al.*, 2016). Feeding *Drosophila* a cocktail of Brefeldin A/Monensin or Tunicamycin decreased the level of invadolysin in male plasma (Figure 3F). On the other hand, treatment with the protein transport inhibitors resulted in no change in the levels of invadolysin in female plasma (Figure 3G). This result suggests that different mechanisms are responsible for the deposition of invadolysin in male vs female hemolymph. In addition, the 46 kDa female form of invadolysin may have a longer half-life than the 111 kDa form observed in males.

### Invadolysin is a component of vertebrate blood

We aimed to determine whether invadolysin was also extracellularly present in higher eukaryotes. Fractionated mouse blood was analysed for the presence of invadolysin. Two different invadolysin antibodies, R2192 and G6456, raised against different epitopes of the protein (Figure 4A) detected an extracellular form of invadolysin (Figure 4B-C). A 53 kDa form of invadolysin was detected in whole mouse blood, and also serum and plasma fractions. The R2192 antibody further detected a 66 kDa form of invadolysin enriched in the blood cell fraction (Figure 4B). An extracellular form of invadolysin at a similar molecular weight of 51-53 kDa was also detected in two samples each of rat and pig plasma (Figure 4D-E).

**Figure 4.**
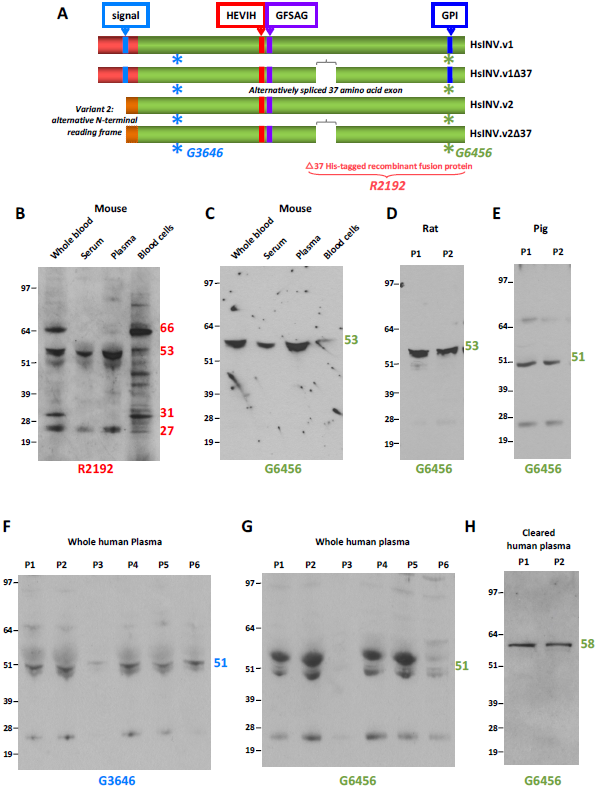
A) Schematic of alternative variants of human invadolysin. Epitopes against which antibodies to invadolysin have been generated are highlighted. B) and C) Mouse blood fractions probed with R2192 and G6456 antibodies identified a secreted form of invadolysin in serum and plasma fractions of mouse blood. R2192 further detected a 66 kDa form of invadolysin that was enriched in the cellular fraction of blood. D) and E) Secreted invadolysin was further identified in rat and pig plasma. F) and G) Human plasma probed using G3646 and G6456 antibodies identified a secreted form of invadolysin. H) Human plasma probed with G6456 anti-invadolysin antibody after abundant proteins have been removed by chromatography. After removal of abundant plasma proteins, the invadolysin signal becomes more prominent and better resolved.

Immunoblotting of human plasma with two antibodies recognising distinct epitopes of invadolysin separated by 408 amino acids, G3646 and G6456, detected invadolysin at ∼51 kDa in six control samples (Figure 4F-G). This band was less readily detected with the G6456 antibody which may be due to the proximity of albumin and immunoglobulin heavy chain, accounting for 70-80% of the total protein content of plasma (Liu *et al.*, 2011; Steel *et al*., 2003). To improve the detection of invadolysin, abundant plasma proteins were removed using a commercially-available abundant plasma protein removal kit (Materials and Methods). Immunoblotting of human plasma by G6456 after depletion of the 12 most abundant proteins dramatically improved detection of invadolysin as evidenced by increased intensity and resolution of invadolysin (Figure 4H). After depletion of the abundant proteins from plasma, invadolysin appears to migrate at 58 kDa rather than ∼51 kDa in whole plasma. This result corroborates the suggested impact on the migration of invadolysin by abundant plasma proteins in the 50-70 kDa molecular weight range. Most importantly, these results demonstrate that deposition of invadolysin into an organism’s circulation is conserved amongst higher eukaryotes.

### Extracellular invadolysin is glycosylated and present in the extracellular vesicular fraction

To simplify the composition of the plasma fraction containing invadolysin, we set about developing a biochemical enrichment for invadolysin. Polyethylene glycol has been used to enrich a particular fraction from a complex protein mixture such as serum or plasma (Haskó *et al.*, 1982). Precipitation of human plasma with different concentrations of PEG (4-10%) clearly shows that invadolysin can be enriched in the PEG-pellet with relatively low concentrations (Figure 5A). We anticipate that the polyethylene glycol precipitation procedure developed in this study will facilitate subsequent detailed analyses of invadolysin structure and function in vertebrates.

**Figure 5.**
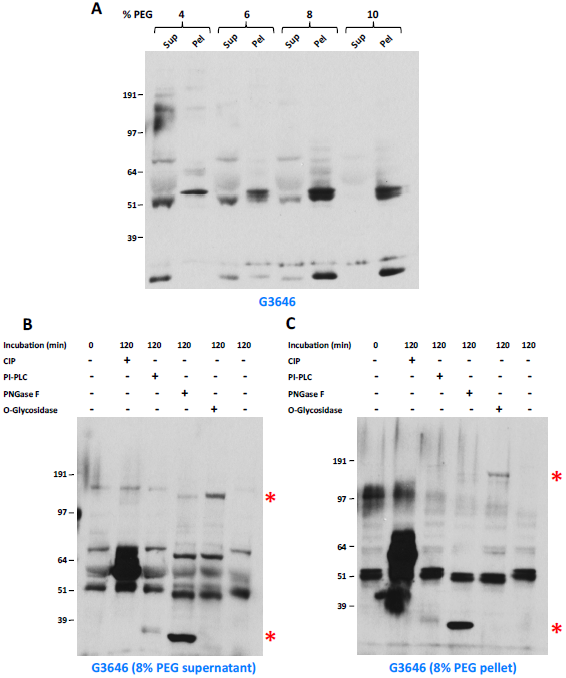
A) PEG 6000 precipitation of human plasma to enrich for invadolysin using PEG concentrations ranging from 4-10%. Aliquots of supernatant and pellet fractions were probed with the G3646 antibody. B) and C) Analysis of post-translational modification of invadolysin in PEG-precipitated supernatant and pellet respectively after 8% PEG 6000 precipitation of human plasma. Samples were treated with CIP/Calf Intestinal Phosphatase (phosphorylation), PI-PLC (GPI-anchor), PNGase F (N-linked glycosylation) and O-Glycosylase (O-linked glycosylation). Enzymatic treatment and subsequent immunoblotting demonstrated that invadolysin in human plasma is N-and O-glycosylated, but likely not GPI-linked. The prominent bands at ∼35 and ∼150 kDa (red asterisks) in the PNGase F and O-Glycosidase lanes represent the added enzymes respectively.

Human plasma invadolysin in the 8% PEG supernatant and pellet and fractions was analysed for N-and O-linked glycosylation, as well as for GPI-anchor addition, following enzymatic treatment and immunoblotting for shifts in electrophoretic migration. PNGase F removes N-linked glycosylation, and O-Glycosidase removes O-linked glycosylation (Magnelli *et al.*, 2011), while PI-PLC is used to remove GPI-anchors (Lehto and Sharom, 2002). While neither calf intestinal phosphatase (CIP) nor PI-PLC treatment resulted in a change to the migration of invadolysin, a faster electrophoretic migration was observed following treatment of both supernatant and pellet fractions with PNGase F and O-Glycosidase (Figure 5B and C). The prominent bands at ∼35 and ∼150 kDa in the PNGase F and O-Glycosidase lanes represent the added enzymes respectively (red asterisks). These results demonstrate that invadolysin in human plasma is N-and O-glycosylated, but not GPI-anchored.

Plasma, as the soluble component of blood, is a complex fraction including numerous different vesicular fractions such as exosomes, microvesicles, membrane particles and apoptotic bodies (EL Andaloussi *et al.*, 2013). Exosomes are cell-derived vesicles that play a vital role in intercellular signaling (Théry *et al.*, 2002). We analysed human plasma fractions enriched for extracellular vesicles for the presence of invadolysin. In this fractionation, the extracellular vesicle fraction contains both exosomes and microvesicles. Purification of this compartment was verified by the presence of flotillin I (Figure 6A). Immunoblotting for invadolysin with the non-overlapping G3646 and G6456 antibodies demonstrated invadolysin to be present in all plasma fractions analysed (Figure 6B and C). However, a strong signal for invadolysin in the plasma fraction enriched for extracellular vesicles, suggests invadolysin may play a role in extracellular vesicle biology.

**Figure 6.**
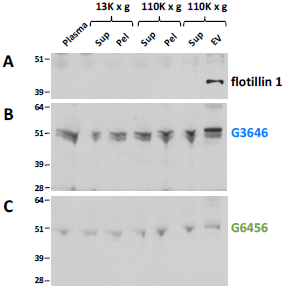
Ultracentrifugation of human plasma to enrich extracellular vesicles and immunoblotting for invadolysin identified invadolysin in the extracellular vesicular fraction. A) Flotillin-1 is utilised as a positive marker for the extracellular vesicle fraction. B) and C) Probing of centrifugation fractions with invadolysin antibodies G3646 and G6456.

## Discussion

The results presented herein are focused on the catalytic motifs and biosynthesis of the conserved metalloprotease invadolysin. Invadolysin has a metalloprotease motif, a third histidine residue and a downstream Met-turn - features characteristic of metzincin metalloproteases (McHugh *et al.*, 2004)(Gomis-Rüth, 2003). Leishmanolysin (Gp63), the closest homolog of invadolysin, is the founding member of the M8 family of metalloproteases (Gomis-Rüth, 2003). Invadolysin however also has a conserved lipase motif downstream of (but very near) the conserved metalloprotease motif. Does invadolysin act as a protease, a lipase or both? No other protein has been shown to have dual proteolytic and lipolytic activity. Our earliest studies demonstrated increased levels of a number of nuclear envelope proteins in *Drosophila* larval extracts and cleavage of lamin by invadolysin in an *in vitro* assay, suggesting the presence of proteolytic activity (McHugh *et al.*, 2004). Invadolysin is also the only protease described as localising to lipid droplets - shown by immunofluorescence as well as biochemical fractionation of cells (Cobbe *et al*., 2009). We showed that invadolysin localises to newly formed lipid droplets in cultured cells following refeeding after serum starvation and increases during adipogenesis of murine 3T3-L1 and human SGBS cells, coincident with an increase in the lipid depot during adipocyte differentiation (Chang *et al*., 2016). These studies thus point toward a role for invadolysin in lipid metabolism and insulin signaling (Chang *et al*., 2016; Cobbe *et al.*, 2009).

Using transgenic *Drosophila* strains that overexpress wild type, protease-or lipase-dead forms of invadolysin, we examined the role of these conserved motifs on lipid and glycogen accumulation. Flies overexpressing wild type or lipase-dead forms of invadolysin achieved significantly higher triglyceride to protein ratios compared to control animals – suggesting the lipase motif is not important in this context. On the other hand, *Drosophila* overexpressing the protease-dead invadolysin transgene were unable to accumulate excess triglyceride, and the triglyceride to protein ratio remained similar to control animals. These results strongly suggest that the protease motif of invadolysin is important for a role in lipid accumulation, though mechanistically how is not clear from these experiments. This is the first direct evidence that proteolytic activity is necessary for invadolysin’s function.

While overexpression of the invadolysin transgenes had no significant impact on the glycogen to protein ratio compared to control animals, in a loss-of-function context, glycogen levels were significantly reduced in *invadolysin* mutants (Chang *et al*., 2016). Impaired insulin signaling in *invadolysin* mutants would be predicted to affect glycogen accumulation, leading to the observed lower glycogen to protein ratio in *invadolysin* mutants (Chang *et al*., 2016). Taken together, our results strongly suggest a role for invadolysin in normal physiology – potentially in an extracellular, endocrine signaling context.

With the goal to understand the biosynthesis of invadolysin, we examined the sequence for potential post-translational modifications. Sequence analysis programmes highlighted an N-terminal signal sequence, numerous N-and O-linked glycosylation sites, and consensus for addition of a C-terminal GPI-anchor. These motifs suggest that variant 1 of invadolysin should be synthesised in the endoplasmic reticulum, while variant 2 may be translated in the cytosol. We thus examined both invertebrate hemolymph and vertebrate blood for the presence of invadolysin. Adult *Drosophila* hemolymph contained invadolysin, but unexpectedly male and female forms differed substantially in molecular weight. Treatment of *Drosophila* with drugs that inhibit protein secretion had a dramatic effect on the invadolysin level in adult male but not in female hemolymph. This observation suggests the testable hypothesis of alternative biosynthetic pathways in males versus females. Whether the expression of sexually dimorphic forms in adult flies is a mere consequence of differentiation, or an active participant in sexual dimorphism remains to be resolved.

Subsequently, we examined plasma from higher vertebrates such as mouse, rat, pig and human to ask whether the secretion of invadolysin was conserved in higher vertebrates. Invadolysin was detected in plasma of all species analysed to date, although we have not yet found any indication for the existence of sexually dimorphic forms in higher organisms. We also identified a distinct form of invadolysin in the cellular fraction of mouse blood. Whether this form is present in particular blood cells is currently under investigation. We determined that human plasma invadolysin was both N-and O-glycosylated, but likely not GPI-linked. Post-translational modification may be important for the interaction with specific binding partners such as regulators or substrates, leading to the modulation of invadolysin’s activity.

What cells or tissues are responsible for the secretion of invadolysin in hemolymph or plasma is currently unclear. Immunoblotting of *Drosophila* male and female gonads for invadolysin identified invadolysin variants similar to those present in hemolymph, suggesting gonads might be responsible for invadolysin production and secretion (or that they take up invadolysin from hemolymph). Many vertebrate plasma proteins are synthesised in the liver, but whether this is true for invadolysin is unknown.

Leishmanolysin/GP63 has been shown to exist in three forms: intracellularly, membrane-anchored at the surface of *Leishmania*, and as a secreted form (Yao *et al*., 2007). Leishmanolysin has also been detected in the exosome-enriched extracellular vesicular fraction. Intriguingly, zymography of exosome fractions from wild type and Leishmanolysin-knockout *Leishmania major* strains showed that knockout of leishmanolysin diminished exosome-associated proteolytic activity (Hassani *et al.*, 2014). We examined fractionated human plasma to determine whether invadolysin is also present in the extracellular vesicular fraction. While clearly present in the purified extracellular vesicle fraction (containing exosomes and microvesicles), invadolysin could also be detected in the other fractions of human plasma. We therefore postulate that invadolysin - like leishmanolysin - is present in different locations, participating in diverse functions.

The soluble fraction of blood is an extraordinarily complex and dynamic assembly of diverse components that plays a wide variety of roles in regulating signal transduction, cell proliferation, differentiation, migration and apoptosis in metazoa (György *et al*., 2011). Our discovery of invadolysin in readily accessible vertebrate plasma opens the doors to biochemical and physiological analyses not particularly tractable with the limited quantities of invertebrate hemolymph obtainable. Identification of proteins interacting with invadolysin will help in understanding the network of invadolysin’s mechanism of action. Is invadolysin active in vertebrate blood? If so, what are its substrates, and how is its activity regulated? Examination of human plasma from control and disease states will shed light on whether invadolysin acts as a biomarker for any pathophysiological states. We anticipate that the study of invadolysin in the extracellular milieu will continue to yield novel insights into this conserved metalloprotease.

## Materials and Methods

### *In silico* and statistical analysis

Amino acid sequences were obtained from the Ensembl genome browser and ClustalX was used for multiple protein sequence alignments. Jalview was used to annotate and highlight key elements of the protein sequence and generate schematics. The programmes used for *in silico* analysis of amino acid sequence features are described in the text. GraphPad Prism 5 was used to carry out statistical analysis of data and generate graphs.

### *Drosophila* experiments

All fly stocks were maintained at 25°C on a standard medium unless otherwise stated. Fly stocks used in this study were: Canton S (wild-type), y w P{nos-phiC31}X; attP40, y w; attP40{UASinv^wt^}/CyO, y w; attP40{UASinv^wt^}/CyO, y w; attP40{UASinv^wt^}/CyO and TubP-Gal4/CyO. For inhibition of the protein secretory pathway, adult flies were fed a cocktail of either 10.6 μM Brefeldin A /2 μM Monensin (Thermo Fisher Scientific) or 60 μM Tunicamycin (Sigma) in 0.15 M sucrose for 7 hours at 25°C.

### Cloning, mutagenesis and generation of transgenics

For generation of transgenic invadolysin constructs, *invadolysin* cDNA (RH66426) was obtained from the *Drosophila* Genomics Resource Center, Indiana University. Transgenic constructs used for generating transgenic flies were made using Thermo Fisher Scientific’s gateway cloning technology. Entry clone was obtained from Invitrogen (now Thermo Fisher Scientific). Expression clones were obtained from the laboratory of Brian McCabe, Columbia University (Wang *et al*., 2012). Mutant versions of the *invadolysin* transgene were generated by site-directed mutagenesis (Carter, 1986). Primers designed for the site-directed mutagenesis reaction were obtained from Sigma and were HPLC-purified. Transgenic constructs were injected into the posterior region of *Drosophila* embryos (where pole cells would form) of a y, w, P{nos-phiC31}X; attP40 *Drosophila* strain. The transgenes were inserted on the second chromosome.

### Total RNA extraction and RT-PCR

Total RNA from *Drosophila* larvae was extracted using the RNeasy Mini Kit (Qiagen) following the manufacturer’s instructions. Total RNA was treated with DNase (Roche) to degrade any genomic DNA contamination. RT-PCR reactions were performed using Superscript III reverse transcriptase (Life Technologies). For specific amplification of transgenic invadolysin, PCR was performed using invadolysin primers (Sigma) designed to selectively amplify the invadolysin transgene. Amplification of transgenic *invadolysin* RNA was performed using GoTaq Green Master Mix (Promega).

### Triglyceride, Glycogen and Protein assay

The samples for triglyceride, glycogen and protein assays were prepared as described previously (Bolukbasi *et al*., 2012). Four adult flies (separated by gender) were homogenised in 400 μl PBS + 0.05% Tween-20. Homogenates were heat-inactivated at 65°C for 5 minutes. Samples were centrifuged at 1200 × g for 1 minute and the supernatant was transferred to a new tube. The supernatant was centrifuged again at 600 × g for 3 minutes. An aliquot of the sample was used for triglyceride, glycogen and protein assays. Triglyceride and glycogen were quantified using BioVision’s Triglyceride/Glycogen Quantification Colorimetric/Fluorometric Kit, and protein was quantified using a Bradford assay kit (Sigma) following manufacturers’ instructions.

### Hemolymph extraction

Hemolymph was extracted from adult flies in EBR solution (130 mM NaCl, 4.7 mM KCl, 1.9 mM CaCl2, and 10 mM HEPES, pH 6.9) containing 20 mM EDTA (Karlsson *et al.*, 2004). Each sample was prepared by pooling hemolymph from 15 adult male or female flies. For collecting hemolymph, the thorax of the fly was punctured and flies were placed in a 0.5 ml tube with a hole in the bottom. The 0.5 ml tube was placed in a 1.5 ml Eppendorf tube containing the 100 μl EBR solution. The tubes were spun for 20 seconds at 300 × g to collect hemolymph. Hemolymph was separated into plasma and cellular fractions by centrifugation for 10 minutes at 300 × g (Karlsson *et al.*, 2004).

### Clean-up of human plasma samples

The Pierce Abundant Protein Depletion Spin Columns (Cat. No.: 85164) from Thermo Fisher Scientific were utilised to remove the 12 most abundant proteins from human plasma. The proteins removed include: α1-Acid Glycoprotein (42 kDa), α1-Antitrypsin (54 kDa), α2-Macroglubulin (85 kDa), Albumin (66.5 kDa), Apolipoprotein A-I (28.3 kDa), Apolipoprotein A-II (17.4 kDa), Fibrinogen (340 kDa), Haptoglobin (18, 45 kDa), IgA (55, 23 kDa), IgG (50, 23 kDa), IgM (65, 23 kDa), and Transferrin (80 kDa).

### SDS-PAGE and immunoblotting

Protein sample preparation and immunoblotting was performed as previously described (Cobbe *et al.*, 2009). Nitrocellulose membranes were probed with primary antibodies listed in the text, and described in previous publications. The Flotillin-1 polyclonal antibody (PA5-19713) was purchased from Thermo Fisher Scientific. Horseradish peroxidase-conjugated secondary antibodies were used, and the immune-signal was detected by ECL (GE Healthcare) with Lumi-Film Chemiluminescent detection film (Roche).

### Enzymatic treatment of plasma

*In silico* predictions for post-translational modification of invadolysin were tested biochemically by enzymatic treatment to remove the predicted modifications. Analysis was done by immunoblotting to observe shifts in protein migration. Human plasma was treated with PNGase F (NEB; P0704S) or a mixture of O-Glycosidase (NEB; P0733S) and *α*2-3,6,8 Neuraminidase (NEB; P0720S) following manufacturer’s instructions. For PI-PLC (Thermo Fisher Scientific), 0.1 enzyme unit was used to treat 10 μg of whole plasma (10 μl).

### Extracellular vesicular fractionation

Freshly collected blood in heparin tubes was centrifuged at 2000 × g for 30 min to obtain plasma. Plasma was then diluted with an equal volume of PBS (136.9 mM NaCl, 2.67 mM KCl, 8.10 mM Na_2_HPO_4_, 1.47 mM KH_2_PO_4_, pH 7.4). Fractionation was then performed at 4°C. Plasma was centrifuged at 13,000 × g for 45 min in Eppendorf tubes. Supernatant was collected and ultra-centrifuged at 110,000 × g for 70 min. The supernatant was discarded and the pellet was resuspended in PBS. Extracellular vesicles were pelleted by ultra-centrifugation at 110,000 × g for 70 min. The pellet containing the extracellular vesicles was resuspended in PBS for subsequent analysis or stored at −80°C.

## Acknowledgements

We would like to thank Dr. Petra zur Lage and Prof. Andrew Jarman (University of Edinburgh) for advice and help with injections of transgenic constructs into *Drosophila* embryos. The McCabe lab (Columbia University) generously donated the *Drosophila* gateway destination plasmid. We would also like to acknowledge both the Bloomington *Drosophila* Stock Centre and FlyBase (a database of *Drosophila* genes and genomes) for their immense contribution to the support of research utilising *Drosophila*. We thank the CIR Blood Resource Centre at the University of Edinburgh for facilitating the examination of control human plasma samples.

